# Extending Prot2Token: Aligning Protein Language Models for Unified and Diverse Protein Prediction Tasks

**DOI:** 10.1101/2025.03.03.641065

**Authors:** Mahdi Pourmirzaei, Ye Han, Farzaneh Esmaili, Mohammadreza Pourmirzaei, Salhuldin Alqarghuli, Kai Chen, Dong Xu

## Abstract

Comprehensive protein function and property prediction remains a major challenge due to the vast diversity of sequences, structural variations, and limited labeled data. Existing models are often specialized to be task-specific, requiring independent training, which limits scalability. To address this, we extend Prot2Token, a unified autoregressive framework that focuses on the post-training alignment of pre-trained protein language models (PLMs), to new applications. Our approach enables next-token prediction across new applications of protein-prediction tasks, including protein-protein structure similarity, 3D structure prediction, mutation stability, post-translational modifications (PTMs), substrate-kinase phosphorylation sites, protein-protein affinity, and protein-ion binding sites. We introduce a self-supervised pre-training stage for the decoder, enhancing model initialization and improving downstream predictions. By integrating a causal autoregressive transformer with a pre-trained ESM-2 encoder, our model effectively aligns diverse protein tasks within a single framework. Additionally, we discuss the opportunities and limitations of this approach, providing insights for future research in optimizing PLMs as a general tool for broader biological applications. Code is available on GitHub Repository.

## 1 Introduction

Proteins are the fundamental building blocks of life, playing a critical role in maintaining human health. However, understanding the complex language of proteins—encoded in their sequences and structures—remains a significant challenge for researchers Shim et al. (2019). This complexity limits our ability to interpret, predict, and design proteins for various biomedical and therapeutic applications.

Protein function prediction is particularly challenging due to the vast diversity of protein sequences, structural variations, and the limited availability of labeled data. Unlike natural languages, protein sequences do not follow explicit syntactic rules understandable by humans, making it difficult for models to learn meaningful representations without extensive biological knowledge Ofer et al. (2021). Protein language models (PLMs) offer a transformative solution by learning meaningful representations of protein sequences, enabling researchers to decode and translate protein data into a more interpretable format An & Weng (2022); Ferruz & Höcker (2022). By leveraging PLMs, we can bridge the gap between raw protein information and human understanding, advancing research in drug discovery, disease mechanisms, and synthetic biology.

While PLMs have significantly advanced protein function prediction, current models require task-specific specialization after pre-training Hu et al. (2023); Roche et al. (2024). This reliance on separate modules for distinct tasks leads to inefficient computational resource use and limited scalability. Most PLMs undergo post-training alignment with specialized architectures for individual tasks, requiring independent training and fine-tuning—an approach that is both time-consuming and resource-intensive Weissenow & Rost (2025). A unified model capable of efficiently handling diverse protein-related tasks would overcome this limitation, streamlining protein function prediction and enhancing its accessibility for real-world applications.

To the best of our knowledge, despite the emergence of foundation models for proteins, no comprehensive framework exists to systematically align them across a broad spectrum of protein prediction tasks. Instead, researchers often modify existing foundation models to suit particular applications Schmirler et al. (2024), such as predicting 3D protein structures from sequences using customized techniques Jumper et al. (2021); Lin et al. (2022). One key limitation is that most existing models are based on BERT-style architectures Unsal et al. (2022), while effective for providing meaningful representation, lack the flexibility needed for diverse and controllable protein generation. In natural language processing (NLP), the transition from BERT-style models to autoregressive GPT-style models has enabled more dynamic and human instruction (prompts) to control the generation process. A similar paradigm shift is necessary in protein research—moving beyond static encoders toward more advanced generative AI approaches that provide more comprehensive predictive capabilities.

Although autoregressive transformer models have been explored for the language of protein—such as ProGen2 Nijkamp et al. (2023), RITA Hesslow et al. (2022), and Ankh Elnaggar et al. (2023)—they struggle with controllability and task alignment. Unlike human language models, which leverage prompt mechanisms for guided generation, protein generative models currently lack robust methods to steer their outputs toward biologically meaningful constraints. This gap hinders their practical application in scenarios requiring fine-grained control over prediction outcomes. Addressing this challenge requires a framework that not only unifies multiple protein-related tasks but also enhances model controllability.

To address these limitations, Prot2Token Pourmirzaei et al. (2024) takes a significant step toward unification of diverse protein-related prediction tasks within a single framework. By introducing an autoregressive interface for existing BERT-style PLMs, it uses next-token prediction for all tasks in an instructive manner through a unified tokenization approach. This design allows PLMs to perform a wide range of predictions. In this paper, we extend and refine Prot2Token to support additional tasks, thereby enhancing its versatility in protein analysis. Specifically, we have adapted it to predict protein-protein structure similarity, 3D structures from sequences, mutation-induced melting temperature changes, six types of post-translational modification (PTM) sites, substrate-kinase phosphorylation sites, protein-protein affinity, and protein-ion binding sites. To facilitate site prediction tasks within the Prot2Token framework, we introduce a self-supervised pre-training stage for the decoder, providing a more effective initialization for downstream predictions. This extension strengthens Prot2Token’s capability as a unified model, reducing the need for task-specific architecture specialization and broadening its applicability in computational biology.

## 2 Related Work

Currently many foundation models exist for proteins Wang et al. (2025), but there is still no general and unified approach to align them for a wide range of protein-prediction tasks. According to our findings, only a few methods adopt general approaches for protein prediction tasks. Prot2Token Pourmirzaei et al. (2024) exemplifies a task-agnostic strategy that employs autoregressive transformers to facilitate alignment in a unified and scalable manner. Another effort, HelixProtX Chen et al. (2024), aims to construct a general model for protein design by integrating various modalities, including text, sequence, and structure; however, this approach remains confined to protein design tasks rather than encompassing broader protein prediction. Additionally, within specialized domains such as PTM, researchers have utilized general models to address entire domains collectively. For instance, PTMGPT2 Shrestha et al. (2024) is a specialized model in the PTMs domain that leverages a pre-trained GPT-2 autoregressive language model to predict multiple PTMs within a single framework.

## 3 Method

### 3.1 Architecture

Our method is based on the Prot2Token framework, incorporating a causal autoregressive transformer, referred to as the decoder, connected to a pre-trained bidirectional transformer, designated as the encoder. Specifically, we initialize the encoder with the pre-trained ESM-2 650M weights Lin et al. (2022), allowing the decoder to access the encoder’s output through cross-attention. To ensure that each task’s unique prediction requirements are met, separate tokenizers and embedding tables are used for the encoder and decoder (see Figure 1). More details about the architecture are presented in Appendix A.1.

**Figure 1:**
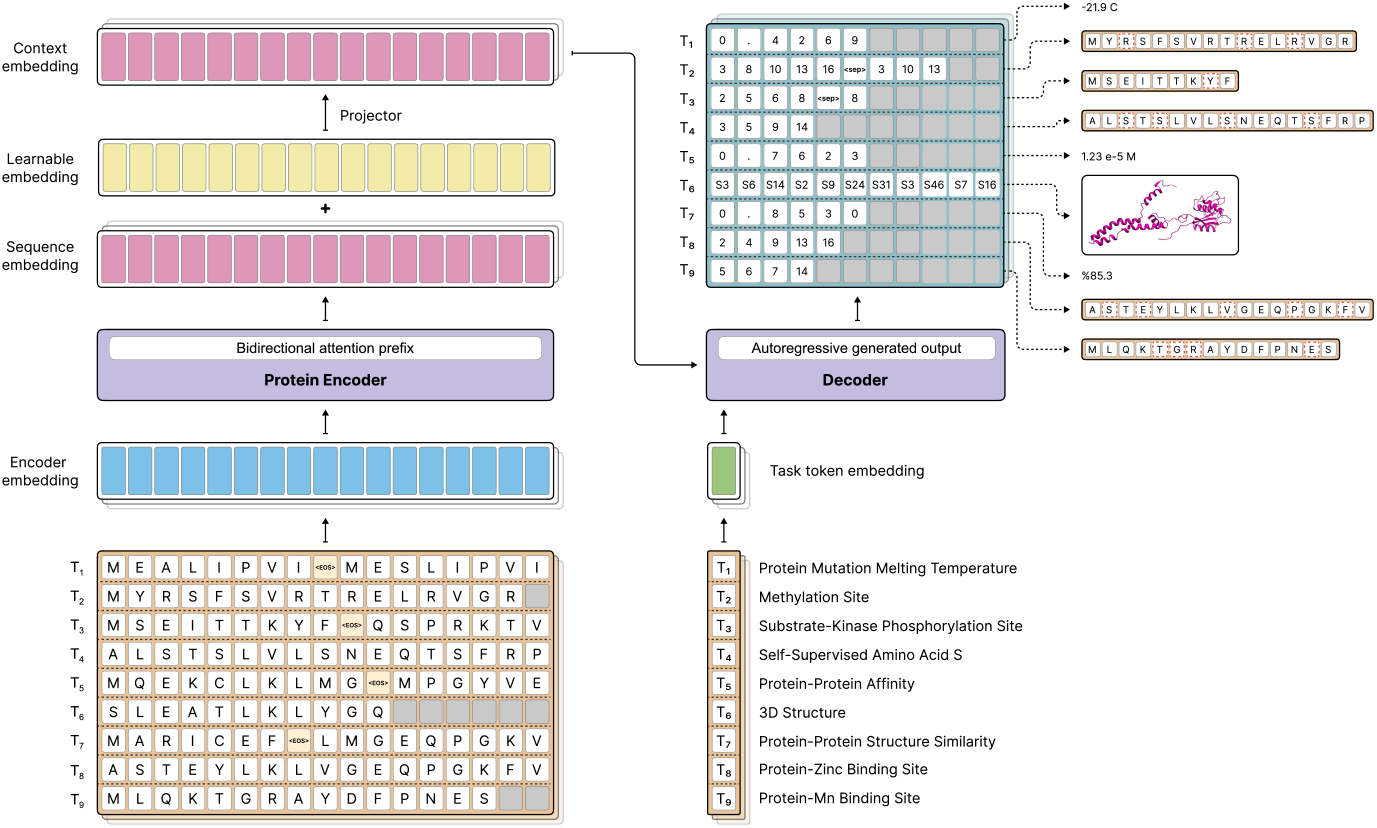
Illustration of the extended Prot2Token framework, which integrates a bidirectional protein encoder with an autoregressive decoder to unify diverse protein prediction tasks. The encoder processes protein sequences with bidirectional attention, generating rich contextual embeddings, while the decoder autoregressively generates structured outputs tailored to specific prediction tasks. The framework supports various tokenization strategies to align sequence-based, structural, and interaction-based protein tasks. On the right side of the figure, predicted tokens are converted to the right format for each task.

### 3.2 Self-Supervised Pre-Training

Unlike the encoder, which can leverage pre-trained weights such as those from ESM-2, the decoder is initialized with random weights in the Prot2token approach. However, prior work in the original Prot2Token paper demonstrated that incorporating self-supervised tasks alongside phosphorylation training can be beneficial for certain tasks. We hypothesize that this advantage arises because the decoder must first grasp the structural patterns of the labels (implicit biases) to generate meaningful predictions. This challenge is particularly pronounced in tasks with larger label vocabularies, such as PTMs, where the available samples may be insufficient for the model to infer these biases effectively, leading to degraded performance.

To mitigate this issue, we introduce a self-supervised pre-training stage that provides the decoder with an initialization before fine-tuning the model on the target task. In this pre-training phase, amino acid sequences serve as inputs, while labels correspond to the positions of specific amino acids. For instance, given a sequence like *“MAGTFAST”*, the target output for a self-supervised task focused on *A* would be the set of positions where it appears in the sequence. These positions are recorded as a sorted set of indices in ascending order, such as {2, 6}. Expanding on this idea, we constructed 20 self-supervised tasks, each dedicated to a different amino acid. A key advantage of these tasks is that they can be generated automatically, eliminating the need for manual annotation. A crucial aspect of these self-supervised tasks is the necessity of freezing the protein encoder. Without this constraint, the model risks collapsing due to shortcut learning, where it exploits spurious correlations rather than learning meaningful representations.

### 3.3 Tokenization of labels

We adopt the tokenization framework from Prot2Token to transform target labels into discrete tokens. Specifically, we apply a regression scheme for protein-protein structure similarity, ProteinProtein Affinity and Protein Mutation Melting Temperature, and use the original Prot2Token PTMs methodology for PTMs and protein-kinase phosphorylation sites tasks in this paper. For protein-ion binding sites, we utilize the same tokenization approach as PTMs but restrict the output tokens to include only the indices of positive binding sites. All potential site tokens and the *<sep>* token are excluded. To handle sequence-to-3D structure mapping, we employ the VQVAE method described in Gaujac et al. (2024). This method converts the backbone coordinates of a PDB file into a sequence of discrete tokens, ensuring that the resulting token sequence matches the length of the corresponding amino acid sequence.

### 3.4 Datasets

This study utilizes a mix of benchmark datasets and custom-curated datasets. PTM prediction, kinase phosphorylation site prediction, protein-ion binding site prediction, protein-protein binding affinity, and 3D structure prediction use datasets we constructed, while other tasks rely on standard benchmarks. For PTM prediction, data from UniProt Consortium (2019) is clustered at 40% similarity (CD-HIT Fu et al. (2012)) and split into training and testing sets, focusing on six key PTM types (Appendix A.2.1). Kinase phosphorylation site data is collected from GPS 6.0 Chen et al. (2023), mapped to UniProt and Kinase.com, clustered at 70% similarity, and split into training, validation, and GPS test sets (Appendix A.2.2).

Protein-ion binding site data is sourced from BioLip2 Zhang et al. (2024a), filtered for proteins with at least 50 residues, clustered at 40% sequence identity, and split accordingly (Appendix A.2.6). Protein-protein affinity data comes from PPB-Affinity Liu et al. (2024), supplemented with missing sequences from RCSB PDB, filtered for single receptor-ligand pairs, and processed with a logarithmic transformation for stability (Appendix A.2.5).

For 3D structure prediction, we use high-confidence UniRef50 Suzek et al. (2015) PDBs from AlphaFold 2 Jumper et al. (2021), filtered by pLDDT scores and tokenized with a 3D structure VQVAE model Gaujac et al. (2024) before splitting into training, validation, and test sets (Appendix A.2.4). Protein-protein structure similarity and melting temperature tasks use ProteinShake Kucera et al. (2024) and ProThermDB/ThermoMutDB Gromiha et al. (2000); Xavier et al. (2021) datasets, respectively (Appendix A.2.3).

## 4. Experiments

We assessed our model across multiple tasks, including protein-protein structure similarity, six PTMs along with protein-kinase phosphorylation site prediction, protein-protein affinity, sequence-to-3D structure mapping, protein-ion binding site identification, and protein mutation melting temperature estimation. For a subset of these tasks, we incorporated a self-supervised pre-training stage for the autoregressive decoder as an initial step. In all experiments, the protein encoder in Prot2Token was initialized using the pre-trained ESM-2 650M model. Optimization was carried out with the AdamW optimizer Loshchilov (2017), applying a weight decay of 0.1 and using beta-1 and beta-2 values of 0.9 and 0.999, respectively, while setting epsilon to 1e-16. The learning rate followed a cosine annealing schedule with an initial warm-up phase Loshchilov & Hutter (2016), starting at 1e-6 and gradually increasing to 5e-5 over the first 256 steps unless stated otherwise. The training was performed using the PyTorch 2 framework Ansel et al. (2024) on a single computational node equipped with four Nvidia A100 GPUs (80GB each).

### 4.1 Protein-Protein Structure Similarity

In our initial experiment, we tokenized ProteinShake protein-protein structure similarity dataset Kucera et al. (2024) and employed the *Structure Split* strategy for evaluation. To ensure consistency during training, we normalized the dataset labels to a range between 0 and 1, maintaining precision up to four decimal places. Each sample comprised two protein sequences, which we concatenated using the *<EOS>* token. The maximum sequence length was set to 1280, and longer sequences were truncated symmetrically to fit within this limit. Additionally, from the total of 33, we finetuned the last four blocks of the protein encoder by unfreezing their weights for training and used batch size of 128 samples per iteration. The results of this experiment are presented in Table 1.

**Table 1:** Structure similarity comparison across different methods. The results are reported on the test set using the Structure Split strategy. All ProteinShake methods rely on 3D structural information. * For the ESM-2 model, a linear layer was added on top of the encoder and it was fine-tuned on the last four blocks of the encoder.

| Method | Prot2Token<br>(Ours) | ESM-2* | ProteinShake<br>(Graph) Kucera et al. (2024) | ProteinShake<br>(Point) Kucera et al. (2024) | ProteinShake<br>(Voxel) Kucera et al. (2024) |
| --- | --- | --- | --- | --- | --- |
| **Spearman R** | 0.5267 | 0.4653 | 0.518 | 0.564 | 0.573 |

### 4.2 Post-Translational Modifications

In the next step, we fine-tuned the model starting from the latest checkpoint obtained during the self-supervised pre-training stage that is reported in Appendix A.3.1. This process involved jointly training six PTMs alongside self-supervised samples. The maximum sequence length for input protein sequences was set to 1024 tokens, and the batch size was adjusted to process 98,304 tokens per iteration.

Notably, while it was possible to exclude self-supervised tasks entirely during fine-tuning, retaining a subset of these samples led to improved generalization and enhanced performance on the protein-kinase phosphorylation site prediction. From the 33 total blocks in the protein encoder, we selectively fine-tuned the last eight blocks by unfreezing their weights for training. The results are presented in Table 2.

**Table 2:**
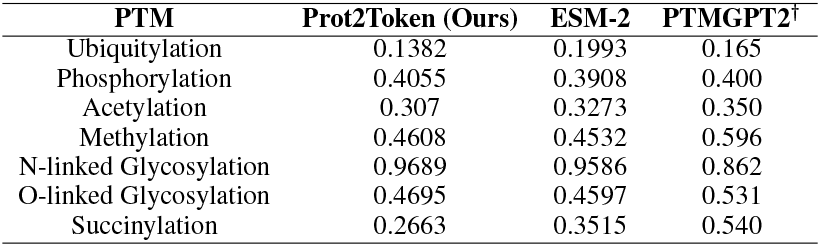
PTMs comparison based on F1 score on our test sets.ESM-2 method is reported in Appendix A.3.2. ^*†*^ there is a strong possibility of <u>data contamination</u> between our test set and the PTMGPT2 training set. As a result, PTMGPT2 may achieve artificially high performance on our test set due to memorization, while its real-world performance on unseen samples could be lower.

#### 4.2.1 Kinase Phosphorylation

Building on the model’s ability to predict PTMs, we extended our approach to include proteinkinase phosphorylation site prediction, a task with significant real-world applications. For this, we selected protein-kinase sequence pairs along with their corresponding phosphorylation sites and jointly trained them alongside 20 self-supervised tasks. The fine-tuning phase started from the latest checkpoint obtained during the self-supervised pre-training stage.

Similar to the PTMs section, in this phase, the self-supervised tasks were reduced to a total of 20,000 samples. Substrate sequences longer than 1,280 amino acids were excluded during training and evaluation. Additionally, the total sequence length, combining substrate and kinase sequences, was capped at 2,048 tokens, with kinase sequences truncated as necessary to fit within this limit. The batch size was set to process 98,304 tokens per iteration. We enabled fine-tuning for the weights of the last eight blocks of the protein encoder.

Table 3, compares our results with two phosphorylation prediction tools, GPS 6.0 and KinasePhos3 Ma et al. (2023). Predictions with scores above 0.7 were classified as true positives. For GPS 6.0, we generated results by selecting each kinase group individually on their platform. Since the training split of GPS 6.0 is not publicly available, there is a risk of <u>data contamination</u> between our validation set and GPS 6.0’s training data. This could result in artificially high-performance estimates for GPS 6.0, potentially reflecting memorization rather than true generalization.

**Table 3:** Comparative results of our method against leading tools (KinasePhos3 and GPS 6.0) across the validation and GPS test.

| Method | Validation Set |  |  | GPS Test Set |  |  |
| --- | --- | --- | --- | --- | --- | --- |
|  | Precision | Recall | F1 | Precision | Recall | F1 |
| KinasePhos3 | **0.9773** | 0.0388 | 0.0747 | 0.0215 | **0.9856** | 0.0421 |
| GPS 6.0 | 0.2323 | 0.4549 | 0.3076 | 0.1564 | 0.5054 | 0.2389 |
| Prot2Token (Ours) | 0.8050 | **0.8200** | **0.8124** | **0.3673** | 0.3103 | **0.3364** |

### 4.3 Protein-Protein Affinity

We applied the same normalization approach on the labels as in the protein-protein structure similarity prediction task, for the protein-protein affinity task. However, before normalization, we transformed the output labels using a logarithmic function, as detailed in Appendix A.1. During training, we initialized the decoder of Prot2Token with randomly assigned weights. We used the same hyperparameters of structure similarity training. The result is presented in Table 4.

**Table 4:** Comparison of protein-protein binding affinity prediction performance between Prot2Token and PPB-Affinity.

| Method | Prot2Token (Ours) | PPB-Affinity<br>Liu et al. (2024) |
| --- | --- | --- |
| **RMSE** | 1.6632 | 2.104 |

### 4.4 3D-Structure Prediction

We trained our model on 8 million randomly selected 3D structures from the training set for 64 epochs, keeping the last 12 blocks of the protein encoder trainable. Throughout training, we monitored model performance on the validation set using the perplexity metric, with results summarized in Table 5. Our analysis of training metrics, combined with computational constraints, suggests that the model is still in the underfitting regime. Extending training on a larger dataset and for a longer duration could further improve validation perplexity. Additionally, by extracting multiple data points from the training process and fitting a regression model, we identified a linear correlation between validation perplexity and the TM-Score of the test set, represented by the equation:

**Table 5:** Performance of the model at the end of training for three different checkpoints, evaluated based on validation perplexity. The table presents the corresponding test set RMSD and TM-score for each checkpoint.

| Valid Set Perplexity | Test Set RMSD | Test Set TM-Score |
| --- | --- | --- |
| 11.53 | 3.89 | 0.7736 |
| 11.41 | 3.79 | 0.7771 |
| 11.08 | 3.72 | 0.781 |

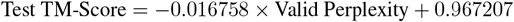

Extrapolating this trend, we estimate that achieving validation perplexity of 6.4, 5.20 and 4.01 would correspond to test set TM-Scores of 0.86, 0.88 and 0.90, respectively. Given the current trajectory, this performance appears feasible with extended training. At the current stage, we evaluated the model’s predictive capability using a checkpoint with a validation perplexity of 11.08 and compared the model’s predictions against AlphaFold 2 predictions at this checkpoint. Failure cases are visualized in Appendix A.3.3.

### 4.5 Protein-ION Binding Site Prediction

For the protein-ion binding site tasks, we focused on four well-known ions, each treated as a separate task and assigned a unique task token. These tasks were jointly trained alongside 20 self-supervised tasks, using the latest checkpoint from the self-supervised pre-training phase as the starting point. During fine-tuning, the number of self-supervised samples was reduced to 50,000. Additionally, protein-ion samples with sequence lengths exceeding 1,280 were excluded, and the batch size was set to 98,304 tokens. Only the last 6 blocks of the encoder (ESM2-650m) were fine-tuned, while all non-encoder parameters of the super model were fully fine-tuned. Notably, while it was possible to omit the self-supervised tasks entirely during fine-tuning, retaining a subset of these samples led to a noticeable improvement in the model’s performance on the supervised protein-ion tasks.

To compare our model’s performance with other available methods, we present the results in Table 6. However, the comparison process was hindered by several challenges which are reported in Appendix A.3.4. Moreover, for all these methods, there is a considerable risk that their training data overlapped with our test sets, potentially biasing the results.

**Table 6:** Comparison of our method’s best performance for each ligand with other available methods on selected ligands based on F1 score. The main values are based on their reported test set performance as described in their respective papers. * Indicates they are reported on our test sets.

| Ligand | Metrics | Prot2Token<br>(Our method) | TargetS<br>Yu et al. (2013) | LMetalSite<br>Yuan et al. (2022) | ZinCap<br>Essien et al. (2019) | MIB2<br>Lu et al. (2022) |
| --- | --- | --- | --- | --- | --- | --- |
| CA<sup>2+</sup> | F1 | 0.6566* | 0.392* | 0.526 (0.7370*) | - | - |
|  | MCC | - | 0.320 (0.431*) | 0.542 (0.7342*) | - | - |
|  | Acc | - | 0.984 (0.977*) | 0.9884* | - | 0.941 |
| MG<sup>2+</sup> | F1 | 0.4603* | 0.433* | 0.367 (0.5560*) | - | - |
|  | MCC | - | 0.383 (0.450*) | 0.419 (0.5773*) | - | - |
|  | Acc | - | 0.990 (0.992*) | 0.9949* | - | 0.946 |
| ZN<sup>2+</sup> | F1 | 0.7594* | 0.660* | 0.76 (0.8299*) | 0.451* | - |
|  | MCC | - | 0.557 (0.660*) | 0.761 (0.8275*) | 0.54 (0.48*) | - |
|  | Acc | - | 0.989 (0.989*) | 0.9953* | 0.870 (0.97*) | 0.948 |
| MN<sup>2+</sup> | F1 | 0.7376* | 0.579* | 0.662 (0.8048*) | - | - |
|  | MCC | - | 0.445 (0.574*) | 0.661 (0.8024*) | - | - |
|  | Acc | - | 0.987 (0.989*) | 0.995* | - | 0.950 |

### 4.6 Protein Mutation Melting Temperature

For this task, we followed the same label preparation strategy as in the Protein-Protein Structure Similarity experiment. The details of the dataset preparation are provided in Appendix A.2.3. Before training, we pre-trained a decoder specifically on the protein-protein structure similarity task while keeping the protein encoder frozen. Then, we initialized the decoder of Prot2Token for this task using that pre-trained decoder. Other than this, we used the same hyperparameters as in the structure similarity training. The results are presented in Table 7.

**Table 7:** Comparison of protein mutation melting temperature prediction performance across different methods.

| Method | Prot2Token (Our method) | GeoDTm-Seq<br>Zhang et al. (2024b) | ESM-2<br>Zhang et al. (2024b) |
| --- | --- | --- | --- |
| RMSE | 8.386 | 8.11 | 7.85 |

## 5 Discussion

The Prot2Token framework was initially designed as a general-purpose approach for unifying diverse protein-related tasks under an autoregressive model. By leveraging a task-agnostic tokenization strategy, it demonstrated the ability to handle multiple protein prediction tasks without requiring task-specific architectural modifications. In this study, we extend Prot2Token to further enhance its applicability by incorporating 3D structure prediction, substrate-kinase phosphorylation site prediction, mutation-induced melting temperature estimation, and protein-ion binding site prediction, broadening its scope beyond the original framework. To improve generalization across these tasks, we introduce a self-supervised pre-training stage for the decoder, ensuring better initialization for site prediction tasks. Our results suggest that this extension enables Prot2Token to align diverse protein-related predictions more effectively.

One key finding is the model’s strong performance in kinase phosphorylation site prediction and protein-ion binding site identification, which achieved competitive or state-of-the-art results. These interactions play a critical role in cellular regulation and drug discovery, making accurate predictions particularly valuable for biomedical research.

For simulating AlphaFold-2 3D structure prediction, Prot2Token demonstrated promising speed improvements, generating predictions nearly 100 times faster than AlphaFold 2 with an average inference time of 1 second on an Nvidia A100 GPU. However, while this efficiency is notable, the accuracy of the generated structures remains lower than that of AlphaFold 2. The current implementation serves as a proof of concept, showing the feasibility of integrating sequence-to-structure prediction within an autoregressive framework. In future work, we will focus on improving accuracy by increasing training data and scaling compute resources, and extending it to predict protein complexes. Given the sign of underfitting observed during training, increasing computational resources by at least 10x could potentially enhance learning capacity and generalization. Additionally, replacing ESM-2 650M with a larger or more advanced PLM could improve the encoder’s representations, leading to better sequence-to-structure mappings.

Despite the advantages of a unified approach, our results highlight challenges, particularly in tasks with limited labeled data. For instance, performance in mutation-induced melting temperature estimation was impacted by the small dataset sizes, reflecting the data dependency of this framework.

In conclusion, this work extends Prot2Token to handle a broader range of protein prediction tasks while maintaining a unified training framework. The model provides flexibility and efficiency, though its performance remains dependent on data availability and task complexity. Moving forward, we plan to further investigate substrate-kinase phosphorylation site prediction and protein-ion binding site identification to refine their predictive capabilities and investigate them deeper. While challenges remain, this study underscores the potential of next token prediction as a step toward general-purpose protein prediction frameworks.

## Meaningfulness Statement

A meaningful representation of life reflects the fundamental principles that govern biological systems, allowing us to interpret the complexity of molecular interactions. Proteins, as essential components of life, encode vast information within their sequences and structures, yet extracting and unifying this knowledge remains a challenge. Our work extends Prot2Token to create a unified framework for protein function prediction, aligning diverse tasks within a single model. By transforming fragmented protein insights into a cohesive representation, we move closer to capturing the underlying order in biological processes, ultimately advancing our ability to model and understand life.

## A Appendix

### A.1 Architecture

The autoregressive transformer models the joint probability of a sequence *x* = (*x*_1_, *x*_2_, …, *x*_*T*_) by decomposing it into conditional probabilities as follows:

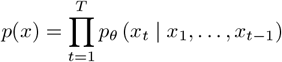

Training is conducted by minimizing the negative log-likelihood of the observed tokens:

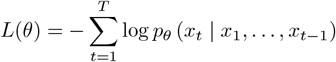

where *θ* represents the model parameters. A causal mask is applied during training to ensure that each token *x*_*t*_ attends only to preceding tokens *x*_1_, …, *x*_*t −*1_. This enforces the autoregressive property, enabling the model to learn contextual representations of the preceding sequence. To refine the standard autoregressive objective, we introduce token-specific weights *w*_*t*_, which allow regulation of the loss contribution from each token. For instance, by setting *w*_1_ = 0, the prompt token (first token) is excluded from the loss computation. For *t* ≥ 2, *w*_*t*_ can be adjusted, enabling non-prompt tokens to have varying importance. The updated training objective becomes:

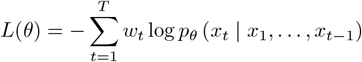

Here, *w*_*t*_ ∈ [0, ∞) is a user-defined parameter that specifies the importance of each token. This approach provides flexibility during fine-tuning by removing the prompt token’s influence on the loss (e.g., assigning it a weight of zero) and focusing on non-prompt tokens. The encoder in our model is identical to the ESM-2 650M architecture. Its output is augmented with a learnable embedding and then reduced from 1,280 to 640 dimensions through a learnable linear projection layer. The decoder consists of a standard causal (autoregressive) Transformer, featuring a hidden size of 640, a feedforward dimension of 2,560, GeLU activations, and 16 attention heads, distributed across 16 blocks. To enhance training efficiency and memory usage, we integrate FlashAttention 2 Dao (2023). To guide the decoder’s predictions for the protein-ion binding site tasks, we introduce task tokens into the process. Unlike the original Prot2Token model, which employed a pre-trained chemical language encoder, we simplify this step by directly mapping each ion type to a specific task token that is fed to the decoder. This strategy allows the model to infer the ion type entirely from the task token, removing the need for a chemical language encoder.

#### A.2 Dataset

##### A.2.1 PTMS

In this section, we describe the process of collecting PTM data. While numerous databases and publications provide PTM data, most only offer sequence fragments, typically 21 amino acids long, with the PTM located at the center position. The largest database with PTM annotations is UniProt, which contains over 200 million protein sequences and provides annotations for more than 200 PTM types and their respective positions for some sequences. We downloaded full-length protein sequences and PTM annotations from UniProt, focusing on annotations in the *Modified Residue, Lipidation, Glycosylation*, and *Cross-link* sections and performed an advanced search in these sections using a wildcard (*) to retrieve all values. This resulted in 106,195 protein sequences from the Reviewed (Swiss-Prot) dataset and 4,173,205 sequences from the Unreviewed (TrEMBL) dataset. To ensure data quality, we exclusively used the protein sequences from the Reviewed (Swiss-Prot) dataset.

We downloaded the 106,195 protein sequences as JSON files for further processing, only sequences with lengths of 1,022 amino acids or fewer were retained. Next, CD-HIT was applied to cluster the sequences based on a similarity threshold of 40% (c = 0.4), grouping sequences with similarity above 40% into the same cluster. Subsequently, we split the data into training and testing sets in a 4:1 ratio, ensuring that sequences within the same cluster were assigned to the same dataset. Given the distribution of PTM types, we focused on six types for this study: Phosphorylation (S), Methylation (R), N-glycosylation (N), O-glycosylation (T), Acetylation (K), and Ubiquitylation (K). Table 8 shows PTM dataset statistics.

**Table 8:** Statistics of PTM datasets.

| PTM type | Annotation in Uniprot | Amino acid | Number of sequences | Number of positions |
| --- | --- | --- | --- | --- |
| Ubiquitylation | Glycyl lysine isopeptide (Lys-Gly) (interchain with G-Cter in ubiquitin) | K | 2,370 | 5,029 |
| Phosphorylation | Phosphoserine | S | 34,260 | 121,398 |
| Acetylation | N6-acetyllysine | K | 9,115 | 23,615 |
| Methylation | Omega-N-methylarginine | R | 1,813 | 3,279 |
| N-linked Glycosylation | N-linked (GlcNAc...) asparagine | N | 30,310 | 11,5767 |
| O-linked Glycosylation | O-linked (GalNAc...) threonine | T | 568 | 2,723 |
| Succinylation | N6-succinyllysine | K | 2,392 | 7,446 |

##### A.2.2 Kinase specific phosphorylation sites

The dataset was gathered from GPS 6.0 and contains 24,160 phosphorylation sites. We mapped IDs from the UniProt database and obtained 13,401 sequences with kinase information. To retrieve kinase sequences, we used Kinase.com and the UniProt database. To reduce sequence similarity, we applied CD-HIT with a 70% similarity threshold to group similar protein substrate sequences.

We kept one representative from each group and selected positive substrate-kinase pairs using two criteria: (1) cross-cluster selection, where pairs from different groups were kept to increase diversity, and (2) within-cluster selection, where only one unique kinase pair per group was retained to avoid repetition. The final dataset includes kinase sequences, kinase information (group/family/kinase), substrate UniProt IDs, substrate sequences, and phosphorylation sites. It contains 386 kinase types across 12 groups. We removed the rare groups *“RGC”* and *“PKL”* due to their poor representation. The dataset was randomly split into training (5,385 unique substrates) and validation (969 unique substrates) sets (Table 9). To test our results against other methods, we used the GPS test set from the *“CMGC”* group, which includes 146 unique substrate-kinase pairs with phosphorylation site information.

**Table 9:** Statistics of kinase phosphorylation site datasets.

| Dataset | Number of samples | Number of p-sites | Number of groups |
| --- | --- | --- | --- |
| All samples | 6,500 | 13,376 | 10 |
| Training set | 5,385 | 10,621 | 10 |
| Validation set | 969 | 2,455 | 9 |
| GPS-test | 146 | 300 | 1 |

##### A.2.3 Protein mutation melting temperature

We used data from GeoStab. The training dataset consists of 4,346 single-point mutations across 349 proteins, sourced from ProThermDB Gromiha et al. (2000) and ThermoMutDB Xavier et al. (2021). The testing dataset contains 571 single-point mutations from 37 proteins, obtained from the same sources.

##### A.2.4 3D-Structure

To construct a high-quality dataset for training our model on sequence-to-structure mapping, we utilized the UniRef50 database, which offers clustered sets of sequences from the UniProt Knowledgebase, reducing redundancy and enhancing computational efficiency. This resource provided us with approximately 67 million unique protein sequences. We then retrieved the corresponding 3D structures for these sequences from the UniProt Predicted Structures Database, which contains models predicted by AlphaFold 2. This effort resulted in the acquisition of 40 million Protein Data Bank (PDB) files. To ensure the reliability of our dataset, we filtered these structures based on their predicted Local Distance Difference Test (pLDDT) scores, a per-residue measure of confidence provided by AlphaFold 2. Structures with a mean pLDDT below 0.85 were excluded, as scores above this threshold indicate high confidence in the predicted local structure. This filtering step reduced our dataset to 11 million PDB files. From this refined collection, we randomly selected two subsets of 2,000 PDB files each, ensuring that all chosen structures had pLDDT scores exceeding 0.90, indicating very high model confidence. These subsets were designated as our validation and test sets, respectively. The remaining structures constituted our training set. Prior to training, all PDB files were converted into discrete tokenized representations using the VQVAE model Gaujac et al. (2024). This process transformed the continuous 3D coordinate data into sequences of discrete tokens, facilitating their use in autoregressive transformers.

##### A.2.5 Protein-Protein Affinity

We used data from PPB-Affinity Liu et al. (2024), the largest publicly available dataset for proteinprotein binding (PPB) affinity. PPB-Affinity provides key information, including crystal structures of protein-protein complexes, PPB affinity values, receptor protein chains, and ligand protein chains. Since PPB-Affinity does not include protein sequences, we retrieved them from the RCSB Protein Data Bank (PDB) based on the protein names provided in PPB-Affinity. To construct a relevant dataset for our model, we applied the following filtering steps:

**1. Chain Filtering** – We removed samples containing more than two chains, retaining only those with a single receptor chain and a single ligand chain.

**2. Mutation Removal** – Samples containing mutated sequences were excluded.

**3. Affinity Label Processing** – For identical protein complexes with multiple PPB affinity values, we averaged the KD (M) values to obtain a single affinity label.

**4. Data Splitting** – The final dataset was split into training (50%), validation (20%), and testing (30%) sets, resulting in 765, 180, and 270 samples, respectively.

The (*KDK*_*D*_) values, representing dissociation constants, were preprocessed to ensure numerical stability and improve model performance. First, a log10 transformation was applied to address the wide dynamic range and skewed distribution of KD values, using the formula: *KD*_log_ = log_10_(*KD* + *ϵ*), where *ϵ* = 10^*−*16^ prevents undefined values for extremely small inputs. The logtransformed values were then normalized to a range between 0 and 1 using Min-Max scaling based on the training dataset’s minimum and maximum *KDlog*_log_ values. Importantly, during model metric calculation and evaluation, both the log-transformation and normalization effects were reversed, ensuring that the calculated metrics accurately reflect the original KD scale. This preprocessing pipeline provided a consistent and interpretable representation of KD values for both model training and evaluation.

##### A.2.6 Protein-Ion Binding Site

We utilized the BioLip2 database to obtain protein interactions with metal ions. BioLip2 primarily relies on the Protein Data Bank (PDB), literature reviews, and other specialized databases. To refine our dataset, we removed DNA/RNA sequences and excluded any protein sequences with fewer than 50 residues. Additionally, we applied CD-HIT with a 40% sequence identity cutoff to cluster the training, validation, and testing datasets. This step ensured a reliable evaluation by maintaining a clear separation between the training and testing datasets. Table 10 provides details on the selected metal ions, including the total number of interacting protein sequences and the corresponding number of residues.

**Table 10:** Protein-ion dataset statistics.

| Chemical Formula | Name | Num Sequences | Binding Sites |
| --- | --- | --- | --- |
| CA<sup>2+</sup> | Calcium Ion | 3043 | 22161 |
| MG<sup>2+</sup> | Magnesium Ion | 2951 | 9494 |
| ZN<sup>2+</sup> | Zinc Ion | 4665 | 23310 |
| MN<sup>2+</sup> | Manganese Ion | 789 | 3315 |

#### A.3 EXPERIMENTS

##### A.3.1 Self-Supervised Pre-training

At the initial stage, we selected 4 million protein sequences from the UniRef50 database Suzek et al. (2015) for training and allocated 4,000 sequences for validation. To expand the dataset, we generated 80 million training samples and 20,000 validation samples by treating each occurrence of an amino acid type within a protein as a distinct training instance. From this pool, we further sampled 1 million training and 1,000 validation samples to construct the final dataset.

For model training, we set the input sequence length to 1,280 and applied a weight decay of 0.01, using a batch size of 192 samples, which corresponds to 73,728 tokens. The training schedule included a warm-up phase of 512 steps. Throughout training, we froze the encoder weights while updating all other parameters. After 16 epochs, the model reached a validation perplexity of 2.31, demonstrating its ability to accurately reconstruct protein sequences from the encoder’s embeddings.

##### A.3.2 PTMs

Training of ESM-2 method was performed for 48 epochs with a cosine annealing learning rate schedule and a warm restart at epoch 24. The initial learning rate is 5 × 10^*−*5^, resetting to 2.5 × 10^*−*5^ at epoch 24, with a minimum learning rate of 0. The AdamW optimizer is used with a weight decay of 1 × 10^*−*2^, and gradient clipping with a norm of 1 is applied to prevent exploding gradients. The dataset is processed with a batch size of 8 and a maximum sequence length of 768 to ensure compatibility with computational constraints.

##### A.3.3 3D structure

During the training process, we encountered multiple interruptions due to various factors, including model collapse, suboptimal learning rates, and unforeseen coding bugs. Given the computational constraints, it was not feasible to maintain a single uninterrupted training session. As a result, we adopted a checkpointing strategy, where training was resumed from the most recent stable checkpoint after each interruption. While this approach allowed us to progress despite hardware limitations, it also introduced challenges related to training continuity, as well as reporting and tracking training logs and metrics. Future iterations of this work would benefit from a more robust computational setup to enable seamless, long-duration training runs.

During inference, we encountered a challenge where the decoder occasionally generated an output sequence with either more or fewer tokens than the actual number of amino acids in the input sequence. To address this issue, we applied a constraint on the end *<EOS>*token probability. Specifically, during inference, we artificially adjusted the probability of the *<EOS>* token, ensuring that it was only allowed if the number of predicted 3D tokens matched the length of the input amino acid sequence. This adjustment effectively enforced sequence alignment and resolved inconsistencies in output length of generated structure. We also demonstrated three failure cases in Figure 4.

**Figure 2:**
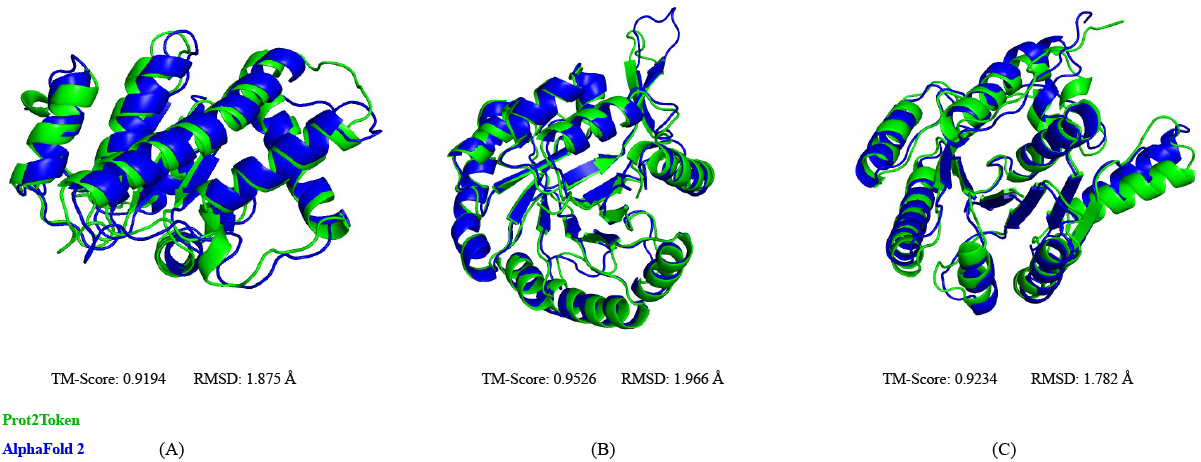
Randomly selected test set samples where our model achieved a TM-score above 0.90. On average, each sample was predicted and converted in approximately 1 second using an Nvidia A100 GPU.

**Figure 3:**
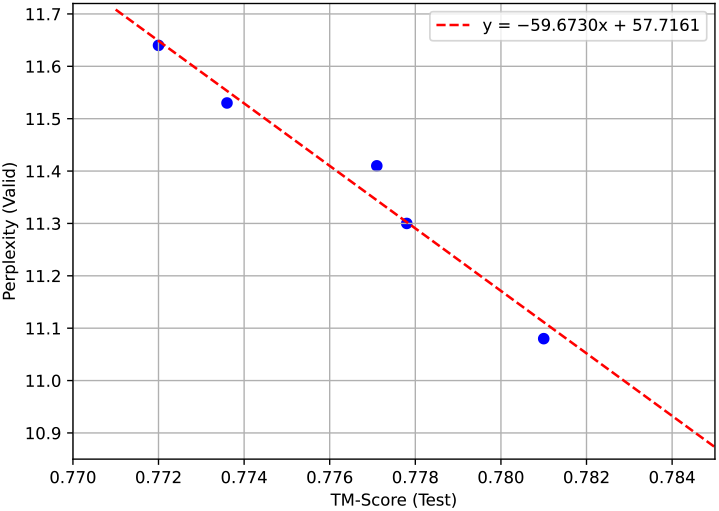
Correlation between validation perplexity and test TM-score. The plot shows a negative linear relationship, with a fitted regression line indicating the estimated trend.

**Figure 4:**
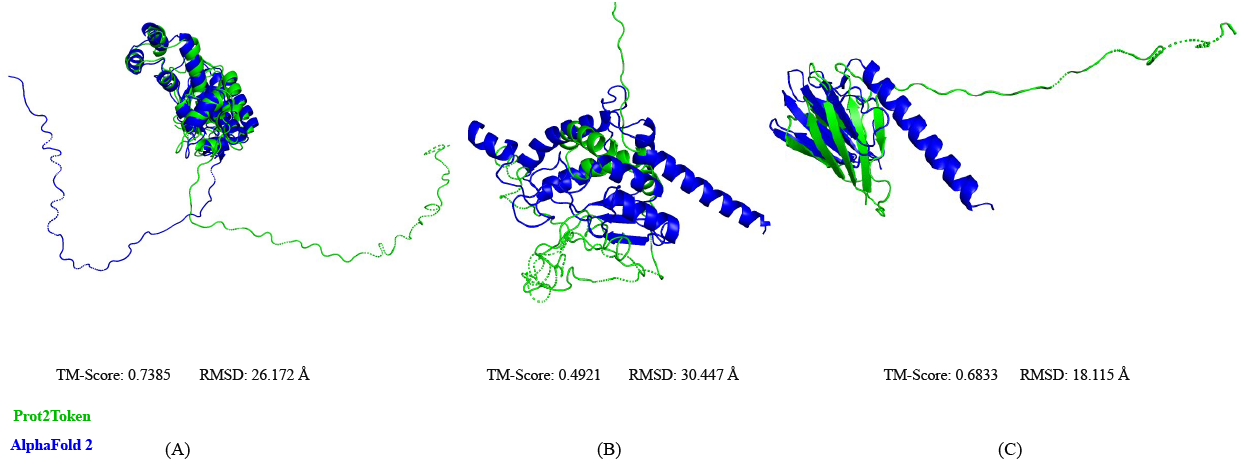
Randomly selected test set samples where the model achieved a TM-score lower than 0.75.

##### A.3.4 Protein-Ion binding site

While comparing our method with other well-known tools, we encountered several challenges. Some web servers were inaccessible during testing, while others only supported single-sample predictions, making bulk evaluations impractical and time-consuming. Specifically, we attempted to evaluate the IonCom Hu et al. (2016) and MIB2 Lu et al. (2022) server tools but faced significant issues: MIB2 exhibited extremely slow response times, and IonCom imposed strict limitations on the number of samples that could be evaluated.

## Notes

https://github.com/mahdip72/prot2token

### Competing Interest Statement

The authors have declared no competing interest.

